# Somatic cells compartmentalise their metabolism to sustain germ cell survival

**DOI:** 10.1101/2025.07.22.666113

**Authors:** Diego Sainz de la Maza, Holly Jefferson, Celine I. Brucker, Sonia Paoli, Marc Amoyel

## Abstract

To ensure success in reproduction, organisms dedicate substantial resources to supporting the germline. In testes, somatic gonadal cells form a barrier that isolates germ cells from circulating nutrients, raising the question of how germ cell metabolism is sustained and how somatic cells ensure sufficient resources are directed to the germline. We use lineage-specific manipulations and metabolite reporters to show *in vivo* that somatic gonadal cells break down circulating sugars to produce and shuttle lactate to the germline, sustaining germ cell survival. We show that somatic cells ensure that carbohydrate metabolism is allocated specifically to germ cell support and that increasing autonomous consumption of carbohydrates in the soma increases germ cell death. Thus, germ cell survival depends on correct metabolic compartmentalisation within gonadal somatic support cells.

## Introduction

Germ cells mediate species survival by producing gametes that enable reproduction. Thus, a considerable proportion of organismal resources is directed towards supporting and nurturing germ cells. Much of this support is provided locally within the gonads by somatic gonadal cells (Messer et al., 2025; O’Donnell et al., 2022). In males, somatic gonadal cells establish a permeability barrier that isolates developing germ cells from blood circulation (Dym and Fawcett, 1970), suggesting that male germ cells depend on nutrients provided locally by the soma. Yet, whether and how somatic gonadal cells provide sustenance to the germline *in vivo* is unresolved.

We use the Drosophila testis as a tractable model to understand how somatic gonadal cells support germline metabolism. Similar to the mammalian testis, germ cells in the fly testis are enclosed by somatic cells called cyst cells (Fig. 1A) (Hardy et al., 1979; Messer et al., 2025). Germline stem cells (GSCs) produce daughters that are enclosed by two somatic cyst cells and subsequently undergo four incomplete and synchronised divisions to form 16 interconnected spermatogonial cells which then mature into meiotic spermatocytes. Around the 4 cell stage, cyst cells establish septate junctions with each other to form a permeability barrier which prevents the developing germ cell syncytium from accessing the external environment (Fairchild et al., 2015). This conservation of male germ cell isolation highlights the importance of tightly controlling the extracellular environment perceived by developing spermatocytes and implies that cyst cells must provide nutrients to sustain germ cell development; yet it remains unknown how they support the germline.

**Figure 1.**
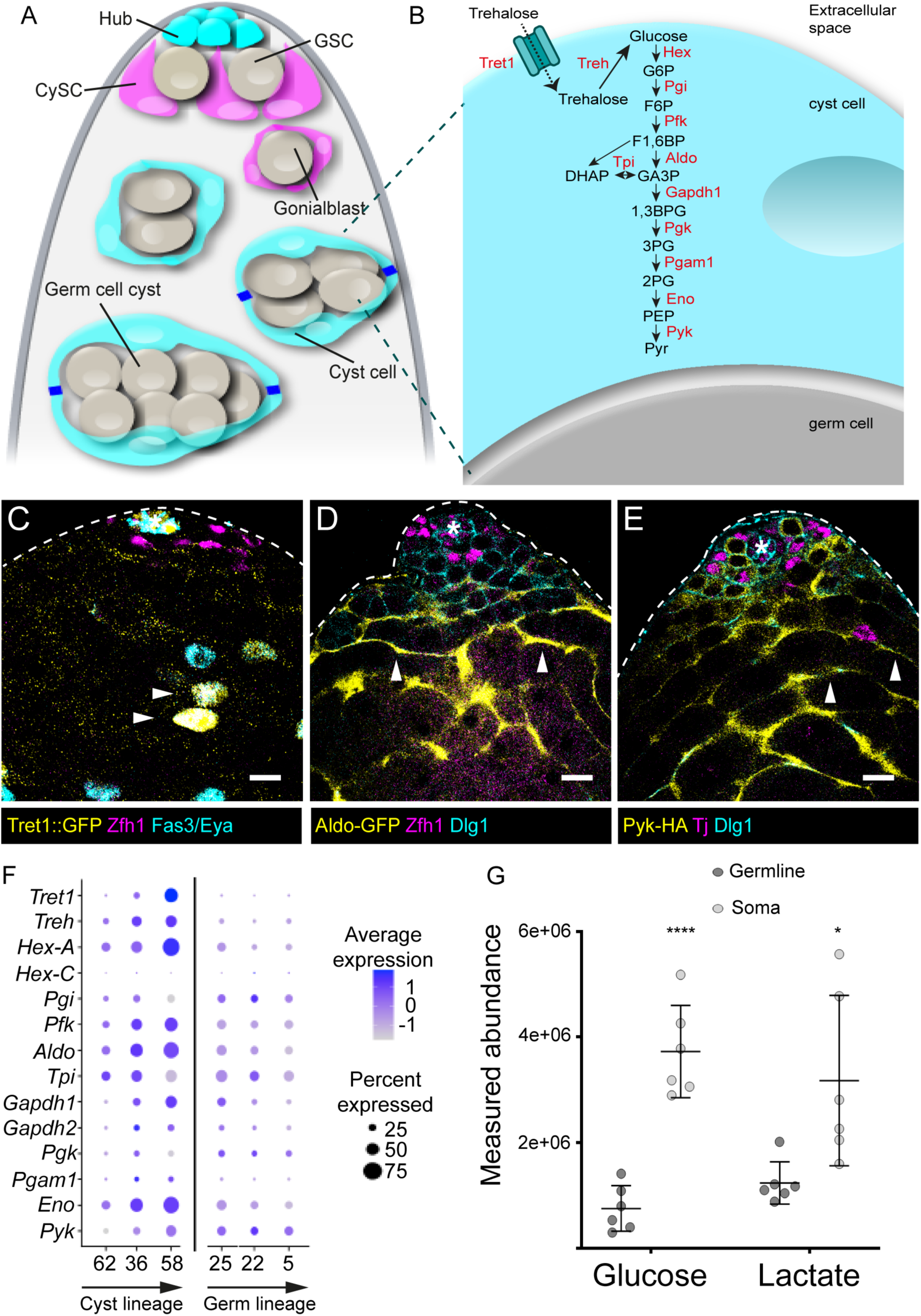
Glycolysis is upregulated in somatic cells. (A) Diagram of the apical tip of a Drosophila testis. The stem cell niche, called hub (cyan), is a group of post-mitotic cells at the tip of the testis (top) which supports two stem cell populations, cyst stem cells (CySCs, magenta) and germline stem cells (GSCs) grey. GSCs divide to give rise to a gonialblast (grey) that is encapsulated by two CySC daughters called cyst cells. The gonialblast divides with incomplete cytokinesis within the envelope formed by the somatic cyst cells (cyan) to form a germ cell cyst of 2, 4, 8 (grey) and 16 (not shown) interconnected cells. Around the 4-cell stage, the two cyst cells form tight junctions (blue), isolating the germline from the external environment. (B) Diagram representing the glycolytic pathway. In flies, the major circulating sugar is trehalose, which is imported into cyst cells by the Trehalose transporter (Tret1) then cleaved into two molecules of glucose. Glucose is then broken down into pyruvate (Pyr) through glycolysis. Enzymes catalysing each step are indicated in red. (C) Confocal image of a testis tip from a *Tret1::nlsGFP* fly showing GFP expression (yellow). Zfh1 (magenta) labels CySCs and their immediate daughters, Fas3 (cyan) labels the hub and Eya (cyan) labels cyst cells. (D) Confocal image of a testis from a fly carrying an *Aldo-GFP* protein trap labelled with GFP (yellow), Zfh1 (magenta) labels CySCs and their immediate daughters and Dlg1 (cyan) labels cell outlines. (E) Confocal image of a testis from a *Pyk-HA* fly showing HA (yellow), Tj (magenta) labels CySCs and early cyst cells, and Dlg1 (cyan) labels cell outlines. All panels: asterisks indicate the hub, arrowheads indicate GFP-expressing cyst cells. Scale bars: 20 μm. (F) Plot showing expression of *Tret1*, *Treh* and glycolytic enzymes according to data from the Fly Cell Atlas in clusters corresponding to CySCs (62), early cyst cells (36 and 58), spermatogonia (25), mid-late proliferating spermatogonia (22) and at the spermatogonium-spermatocyte transition (5). The size of the dots indicates the percent of expressing cells in each cluster, and the colour represents the expression level. (G) Measured abundance of glucose and lactate obtained by LC-MS in sorted somatic and germ cells. **** denotes P<0.0001, * denotes P < 0.05, student’s t test.

Here we demonstrate metabolite exchange between somatic gonadal cells and germ cells *in vivo*. We show that lactate, a product of glycolysis in cyst cells, is shuttled to the germline to support its survival. Furthermore, we show that cyst cells compartmentalise their metabolism, ensuring that carbohydrates, through glycolysis, are dedicated to germline support, and that disrupting this allocation of resources in cyst cells impacts germ cell survival. Our work thus highlights the importance of somatic gonadal cells in ensuring that sufficient resources reach the germline, and demonstrates that metabolic compartmentalisation within cyst cells is crucial to their support function within the gonads.

## Results

### Glycolysis is upregulated in differentiated somatic cells

Cyst cells are produced by cyst stem cells (CySCs), differentiating when two cyst cells enclose a germ cell (Gonczy and DiNardo, 1996; Hardy et al., 1979). We previously showed that differentiating cyst cells upregulate genes encoding many metabolic enzymes compared to CySCs (Sainz de la Maza et al., 2022). We noted that many genes encoding glycolytic enzymes (see diagram in Fig. 1B) were represented within the Gene Ontology (GO) terms most significantly enriched in differentiating cyst cells. In particular, *Hexokinase-A (Hex-A), Phosphoglucose isomerase (Pgi), Phosphofructokinase (Pfk), Triose phosphate isomerase (Tpi), Glyceraldehyde-3-phosphate dehydrogenase 1 (Gapdh1), Enolase (Eno)* and *Pyruvate kinase (Pyk)* were upregulated more than 2-fold in cyst cells relative to CySCs (Sainz de la Maza et al., 2022). Thus, we hypothesised that glycolysis is upregulated during cyst cell differentiation. We validated these findings by examining expression reporters for key enzymes. In Drosophila, the main circulating sugar is trehalose, which is imported into cells by the Trehalose transporter 1 (Tret1, also known as Tret1-1) (Fig. 1B) (Kanamori et al., 2010). A transgene expressing nuclear GFP under the control of the *Tret1* enhancer (Hertenstein et al., 2021) showed GFP expression in the testis, specifically in hub cells and in Eya-positive differentiating cyst cells (Fig. 1C), but was absent from Zfh1-positive CySCs and Vasa-positive germ cells. Similarly, a protein trap GFP fusion with Aldolase 1 (Aldo), the enzyme that splits the 6-carbon fructose 1,6 bisphosphate into two 3-carbon sugars (Fig. 1B), was expressed highly in the cytoplasm of Eya-positive differentiated cyst cells and more weakly in Zfh1-positive CySCs (Fig. 1D). Finally, we examined the expression of a tagged form of Pyruvate kinase (Pyk), which catalyses the final step in glycolysis, forming pyruvate and ATP. Pyk-HA was expressed in GSCs and early germ cells, but expression decreased in later stages of germ cell differentiation (Fig. 1E). By contrast, in the somatic lineage, HA staining was low in CySCs adjacent to the hub and increased in intensity as cyst cells differentiated. Analysis of the recent Fly Cell Atlas testis dataset of single-cell gene expression (Li et al., 2022; Raz et al., 2023) revealed a similar pattern. We compared gene expression of the clusters corresponding to spermatogonia and the cyst cells associated with spermatogonial stages (Fig. S1A-C). GO analysis of the differentially expressed genes revealed that amino acid metabolism and carbon metabolism were enriched in cyst cells, but examination of the genes within these categories revealed overlapping genes including genes of the glycolysis pathway (Fig. S1D). Plotting expression of glycolytic genes showed that these were most highly expressed in differentiating somatic cells, increasing as cyst cells differentiated, while in germ cells, expression decreased during differentiation (Fig. 1F). Together, these data show that glycolytic gene expression increases in cyst cells as they differentiate, and suggest that cyst cells have higher glycolytic activity than germ cells.

To test whether glycolytic pathway activity was indeed higher in somatic cells, we used liquid chromatography mass spectrometry to compare levels of glucose and lactate between cyst and germ cells, which we sorted by FACS using *traffic jam (tj)-Gal4* to drive GFP expression in somatic cyst cells. Analysis of the mass spectrometry data by principal component analysis revealed that germ cells and cyst cells segregated reproducibly, suggesting different metabolic profiles (Fig. S1E) and indeed, we identified thousands of annotated peaks as differentially enriched (Fig. S1F). Importantly, somatic cells had significantly higher levels of both glucose and lactate than germ cells (Fig. 1G). Altogether, these data show that cyst cells have higher levels of glycolytic gene expression than germ cells, as well as higher amounts of glucose and of the glycolytic product lactate. Notably, the increase in gene expression occurs around the time or soon after the two cyst cells encapsulating a developing germ cell cyst form tight junctions, isolating the germline (Fairchild et al., 2015). Based on these observations, we hypothesised that cyst cells may upregulate glycolysis to provide the developing germline with glycolytic byproducts and sustain germ cell development.

### Glycolytic enzymes in cyst cells non-autonomously maintain germ cell survival

To test this hypothesis, we sought to knock down glycolytic enzymes in somatic cyst cells and assess the effect of the knockdown on the germline. We expressed RNAi constructs that we previously validated for effectiveness (see Table S1) in the cyst lineage (Fig. S2A), using *tj-Gal4* together with *hh-Gal80* to prevent Gal4 activity in hub cells (Herrera et al., 2021). Knockdown of enzymes involved in trehalose import or processing, or in glycolysis resulted in dying germ cells, as evident by the presence of rounded germ cell cysts, positive for the acidophilic dye Lysotracker (Fig. 2A,B) (Yacobi-Sharon et al., 2013). Quantification of dying germ cell cysts per testis revealed a significant increase in most glycolytic knockdowns (*Tret1, Trehalase (Treh), Pfk, Gapdh1, Eno, Pyk*) compared to control (Fig. 2C) and a trend towards increased numbers in others (*Pgi* and *Phosphoglycerate mutase 1 (Pgam1)*). Only *Phosphoglycerate kinase* (*Pgk)* knockdown appeared unchanged compared to control and this RNAi was an outlier in subsequent experiments, suggesting that this line may not effectively target *pgk*. Importantly, these knockdowns did not result in any noticeable changes in cell type composition in the testis: CySCs, identified as Zfh1-positive, Eya-negative cells were still present, as were Eya-positive cyst cells (Fig. 2A,B). Germ cell morphology, as assessed by Vasa staining, also appeared grossly normal with all stages present from germline stem cells (GSCs) adjacent to the hub to spermatogonial and spermatocyte cysts.

**Figure 2.**
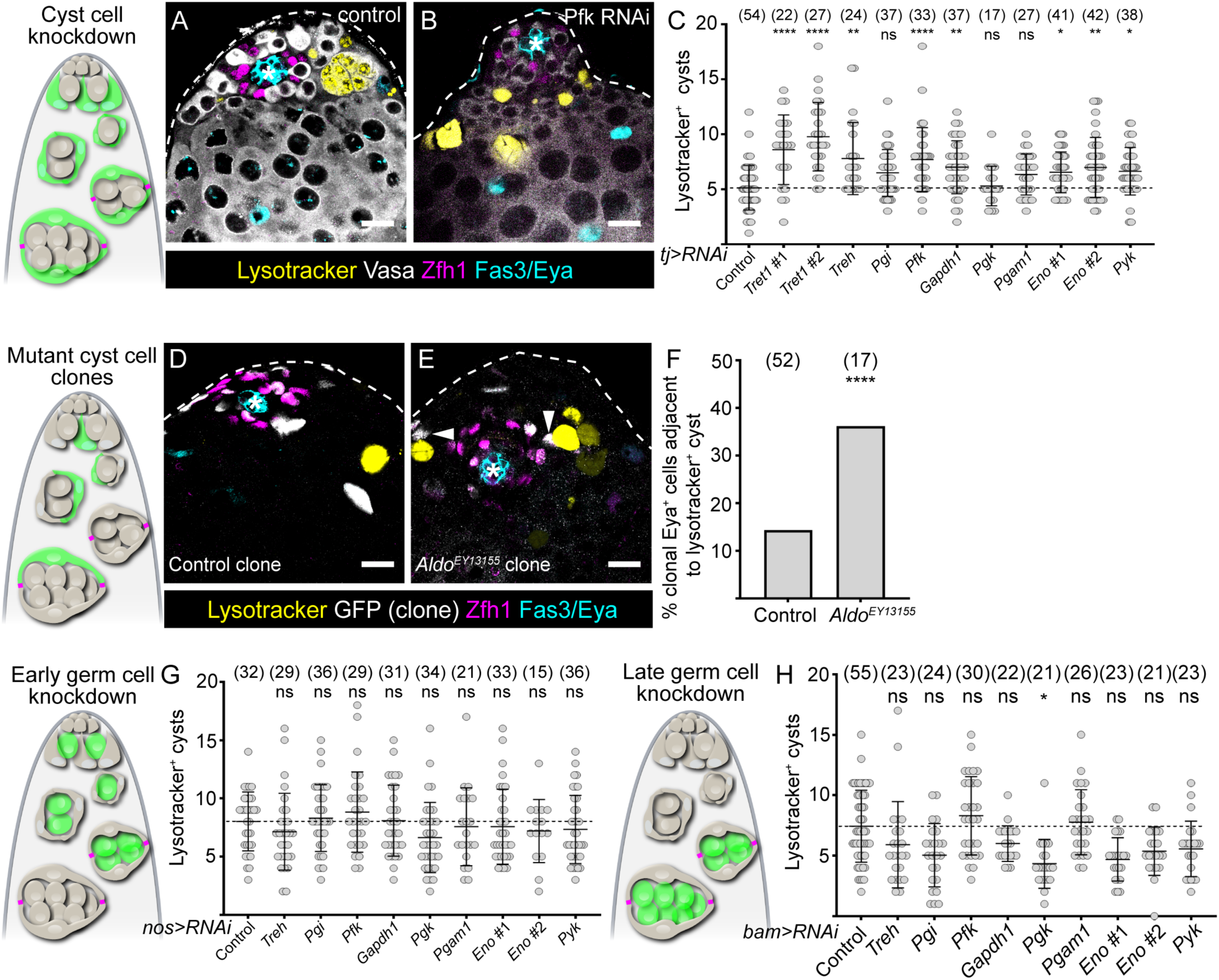
Somatic glycolysis is required for germ cell survival. Diagrams show testis apical tips and highlight the lineage that is manipulated in green. (A,B) Confocal images of control testes (A) and testes in which *Pfk* was knocked down in cyst cells with *tj-Gal4* (B) labelled with Lysotracker (yellow) to label dying germ cell cysts, and antibodies against Vasa to label the germline (white), Zfh1 (magenta) to label CySCs and their immediate daughters, Fas3 and Eya (cyan) to label the hub and cyst cells, respectively. (C) Graph showing the number of Lysotracker-positive cysts per testis upon knockdown of glycolytic enzymes in cyst cells with *tj-Gal4*. Statistical significance was assessed using the Kruskal-Wallis test followed by Dunn’s multiple comparisons, relative to the control. (D,E) Images of testes containing positively-marked control (D) and A*ldo^EY13155^* (E) somatic clones. Lysotracker (yellow) labels dying germ cell cysts, GFP (white) labels clonal cells, Zfh1 (magenta) labels CySCs and their immediate daughters, Fas3 and Eya (cyan) label the hub and cyst cells, respectively. Arrowheads in E indicate clonal cyst cells adjacent to Lysotracker-positive germ cell cysts. (F) Graph showing the ratio of clonal Eya-positive cells adjacent to Lysotracker-positive cysts for control and A*ldo^EY13155^* clones. **** indicates P < 0.0001, Fisher’s exact test. (G,H) Graphs showing the number of Lysotracker-positive cysts per testis upon knockdown of glycolytic enzymes in early germ cells with *nos-Gal4* (G) and late germ cells with *bam-Gal4* (H). Statistical significance was assessed using the Kruskal-Wallis test followed by Dunn’s multiple comparisons, relative to the control. **** denotes P<0.0001, ** denotes P < 0.01, * denotes P < 0.05. All graphs: N values are shown in brackets and refer to the number of testes analysed. Scale bars: 20 μm.

We showed that disrupting trehalose or glucose metabolism did not impact the self-renewal of CySCs or survival of cyst cells: we observed no significant changes in the number of CySCs (Zfh1-positive, Eya-negative cells) or of cyst cells staining positive for an antibody against the activated form of the effector caspase Death Caspase-1 (Dcp-1) upon knockdown of this pathway (Fig. S2B,C). To validate these results independently, we generated CySCs and cyst cells homozygous mutant for an allele of *Aldo*, for which we did not have a functional RNAi. Lysotracker-positive germ cells were found adjacent to *Aldo* mutant cyst cells with a higher frequency compared to control clones (Fig. 2D,E, arrowheads; 36% compared to 14% in controls, P < 0.0001, Fisher’s exact test, Fig. 2F), indicating that *Aldo* loss in cyst cells non-autonomously reduces germline survival. *Aldo* mutant clones appeared indistinguishable from control clones, with several CySCs labelled, suggesting that the labelled cells were capable of proliferating and of producing other CySCs, despite a small but not significant reduction in *Aldo* mutant clone recovery compared to controls (Fig. S2D). Altogether, these data indicate that disrupting trehalose uptake or catabolism into pyruvate through glycolysis does not discernibly impact self-renewal, differentiation or survival of the cyst lineage, but non-autonomously results in germ cell death.

Next, we asked whether glycolytic enzymes were required autonomously in germ cells for their survival. We used *nanos-Gal4* to drive expression in GSCs and early spermatogonia (Fig. S3A) and *bam-Gal4* to drive expression in late spermatogonia and spermatocytes (Fig. S3B). Knocking down glycolytic enzymes with either driver did not result in any increase in the number of Lysotracker-positive dying cysts (Fig. 2G,H). We confirmed that germline knockdown was effective by observing decreased expression of the Pyk-HA fusion protein with an anti-HA antibody upon knockdown of *pyk* with *nanos-Gal4* (Fig. S3C,D). In sum, although knockdown of genes encoding glycolytic enzymes in the soma results in germ cell death, these genes appear to be dispensable in the germline itself, suggesting that germ cells depend on glycolysis in cyst cells to provide them with a critical metabolite, or metabolites, that they need to survive.

### Cyst cell-derived lactate is required for germ cell survival

Our results indicate that enzymes across the whole glycolytic pathway are required for germ cell survival, suggesting that the metabolite(s) produced in cyst cells that support the germline are likely pyruvate or a product of pyruvate metabolism. Cells can either transform pyruvate into lactate through the action of Lactate dehydrogenase (Ldh), or import pyruvate into mitochondria (Fig. 3A). We first tested whether lactate production was required for germ cell survival by knocking down *ldh* in cyst cells. We used a genetically-encoded single fluorophore reporter for lactate, CanlonicSF, to measure lactate levels in cyst cells (Aburto et al., 2022). As expected, *ldh* knockdown resulted in significantly lower fluorescence of CanlonicSF compared to control (Fig. 3B-D). We measured germ cell death and found that somatic *ldh* knockdown resulted in a significant increase in the number of Lysotracker-positive dying germ cell cysts (Fig. 3E). Increased germ cell death upon *ldh* knockdown was a non-autonomous effect and not due to disruptions to cyst cell development as neither CySC numbers nor proliferation were affected (Fig. S4A,B). Next, we asked whether somatic *ldh* knockdown could induce germ cell death indirectly, since cyst cell death can induce germ cell death in starved animals (Yang and Yamashita, 2015). We did not observe an increase in Dcp-1 positive cells upon *ldh* knockdown (Fig. S4C), suggesting that *ldh* is not necessary for cyst cell survival. To further validate that cyst cell death did not precede germ cell death, we inhibited apoptosis autonomously in cyst cells by over-expressing Death-associated inhibitor of apoptosis 1 (Diap1). Importantly, Diap1 expression did not prevent the increase in Lysotracker-positive cysts caused by *ldh* knockdown in cyst cells (Fig. S4D), indicating that germ cells do not die as a consequence of apoptosis in cyst cells.

**Figure 3.**
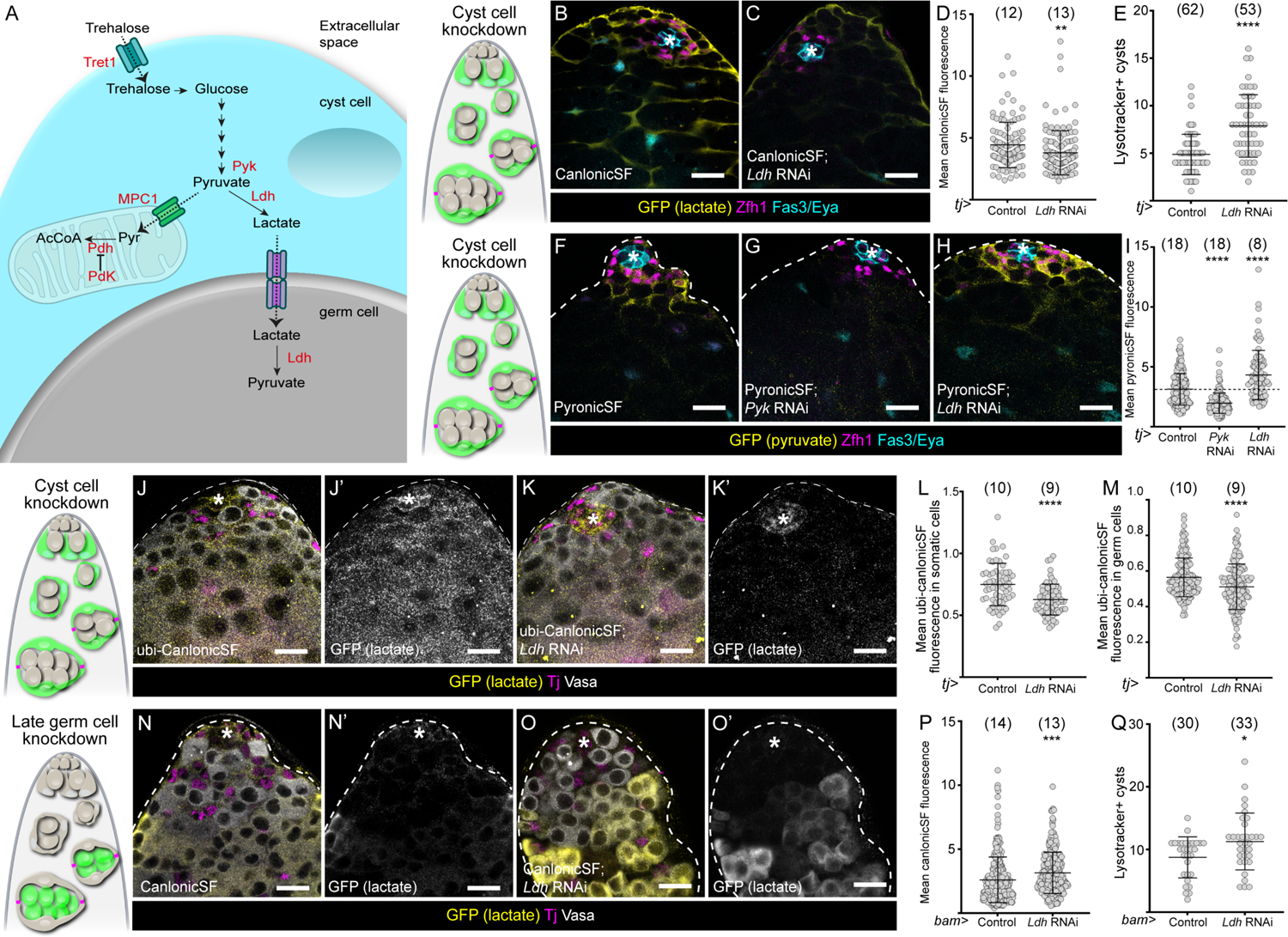
Cyst cell-derived lactate is consumed by germ cells. (A) Diagram showing the possible uses of glycolysis-derived pyruvate in cyst cells. Lactate dehydrogenase (Ldh) catalyses the transformation of pyruvate into lactate, which can be exported via a monocarboxylate transporter. Ldh can also catalyse the reverse reaction once lactate is imported into the germline. Alternatively, pyruvate can be imported into mitochondria via the mitochondrial pyruvate carrier (Mpc1), and decarboxylated into acetyl-CoA (AcCoA) by the Pyruvate dehydrogenase complex (Pdh). Activity of Pdh is inhibited by the Pyruvate dehydrogenase kinase (Pdk). (B,C) Confocal images of testes in which the lactate reporter CanlonicSF (yellow) was expressed in cyst cells with *tj-Gal4* in control testes (B) or together with *Ldh* knockdown (C). Zfh1 (magenta) labels CySCs and their immediate daughters, Fas3 and Eya (cyan) label the hub and cyst cells, respectively. (D) Graph showing the mean intensity of CanlonicSF fluorescence in cyst cells in control and testes in which *Ldh* was knocked down in cyst cells. ** denotes P < 0.01, student’s t test. (E) Graph showing the number of Lysotracker-positive germ cell cysts in control testes and testes in which *Ldh* was knocked down in cyst cells. **** denotes P < 0.0001, Mann-Whitney test. (F-H) Confocal images of testes in which the pyruvate reporter PyronicSF (yellow) was expressed in cyst cells with *tj-Gal4* in control (F) or together with *Pyk* (G) or *Ldh* knockdown (H). Zfh1 (magenta) labels CySCs and their immediate daughters, Fas3 and Eya (cyan) label the hub and cyst cells, respectively. (I) Graph showing the mean intensity of PyronicSF fluorescence in cyst cells in control testes and testes in which *Pyk* or *Ldh* were knocked down in cyst cells. **** denotes P < 0.0001, Kruskal Wallis and Dunn’s multiple comparisons tests. (J, K) Confocal images of testes in which the lactate reporter CanlonicSF (yellow) was expressed ubiquitously in control testes (J) or testes in which *Ldh* was knocked down in cyst cells. (K). Tj (magenta) labels CySCs and early cyst cells, Vasa (white) labels germ cells. (L) Graph showing the mean intensity of CanlonicSF fluorescence in cyst cells in control testes and testes in which *Ldh* was knocked down in cyst cells. **** denotes P < 0.0001, student’s t test. (M) Graph showing the mean intensity of CanlonicSF fluorescence in germ cells in control testes and testes in which *Ldh* was knocked down in cyst cells. **** denotes P < 0.0001, student’s t test. (N,O) Confocal images of testes in which the lactate reporter CanlonicSF (yellow) was expressed in germ cells with *bam-Gal4* in control (N) or together with *Ldh* knockdown (O). Tj (magenta) labels CySCs and early CCs, Vasa labels the germline (white). (P) Graph showing the mean intensity of CanlonicSF fluorescence in germ cells in control and *Ldh* knockdowns. *** denotes P < 0.001, student’s t test. (Q) Graph showing the number of Lysotracker-positive cysts in control testes and testes in which *Ldh* was knocked down in germ cells. * denotes P < 0.05, Mann-Whitney test. All graphs: N values are shown in brackets and refer to the number of testes analysed. Scale bars: 20 μm. Asterisks indicate the hub. Diagrams to the left represent testis apical tips and highlight the lineage where gene expression was manipulated in green.

Altogether, these results indicate that lactate production is necessary in cyst cells to non-autonomously prevent germ cell death, suggesting that lactate supports germ cell survival. However, knockdown of *ldh* may affect flux through glycolysis, such that other glycolytic metabolites would also be reduced by this manipulation (Hosios and Vander Heiden, 2018); therefore we sought to determine if *ldh* knockdown reduced glycolytic products. We examined pyruvate levels using PyronicSF (Arce-Molina et al., 2020). As a control, we knocked down *pyk*, the product of which catalyses pyruvate production, and observed a decrease in fluorescence (Fig. 3F,G). By contrast, knockdown of *ldh* resulted in increased fluorescence (Fig. 3F,H,I), suggesting that pyruvate accumulates in cyst cells. Since *pyk* and *ldh* knockdown both result in decreased lactate production and increased germ cell death but have opposite effects on pyruvate accumulation, it is likely that lactate, and not another glycolytic intermediate, is the key metabolite that supports germ cell survival.

Next, we expressed CanlonicSF under the control of a ubiquitous promoter to ask whether knocking down *ldh* in cyst cells non-autonomously affected lactate levels in the germline. Somatic knockdown of *ldh* resulted in decreased fluorescence, both autonomously in cyst cells, as above, and non-autonomously in germ cells (Fig. 3J-M). This observation supports the idea that lactate is transferred from cyst cells to germ cells. If this is true, we hypothesised that to be utilized in germ cells, somatic-derived lactate would be transformed back into pyruvate by Ldh (Fig. 3A), which can catalyse both reactions. Indeed, knockdown of *ldh* in germ cells using *bam-Gal4* resulted in increased CanlonicSF fluorescence (Fig. 3N-P). Moreover, unlike in the case of glycolytic enzymes (Fig. 2H), *ldh* knockdown in germ cells resulted in increased numbers of Lysotracker-positive cysts (Fig. 3Q), indicating that Ldh-dependent conversion of lactate into pyruvate sustains germ cell survival. Altogether, our results show that glycolysis in cyst cells fuels the production of lactate, which is transferred to germ cells which consume lactate but do not produce it.

### An uncharacterised monocarboxylate transporter mediates lactate transport

Since our results indicated that lactate was transferred from cyst cells to the germline, we asked whether monocarboxylate transporters (MCTs), which mediate transport of metabolites across cell membranes, were required for this transport. We focused on the Solute Carrier 16 (SLC16) family of MCTs, which has 16 members https://flybase.org/reports/FBgg0000669 (Halestrap, 2013). Using the Fly Cell Atlas, we determined that 3 genes encoding MCTs were highly expressed in cyst cells: *CG8034, CG13907* and *hermes* (*hrm*) (Fig. S5A). We knocked down each of these in cyst cells and found that *CG8034* knockdown resulted in increased numbers of Lysotracker-positive germ cell cysts (Fig 4A). To test whether the product of *CG8034* could indeed transport lactate, we examined fluorescence of the CanlonicSF reporter, and found that knockdown of *CG8034* resulted in accumulation of lactate in cyst cells (Fig. 4B-D). Next, we used the ubiquitously-expressed CanlonicSF to ask whether knockdown of *CG8034* in cyst cells affected lactate levels in the germline and found that CanlonicSF fluorescence was significantly decreased in germ cells and concomitantly significantly increased in cyst cells (Fig. 4E-H), consistent with reduced transport of lactate from cyst cells to the germline. Due to its role in delivering lactate to germ cells, we renamed *CG8034, milkman (mlm)*.

**Figure 4.**
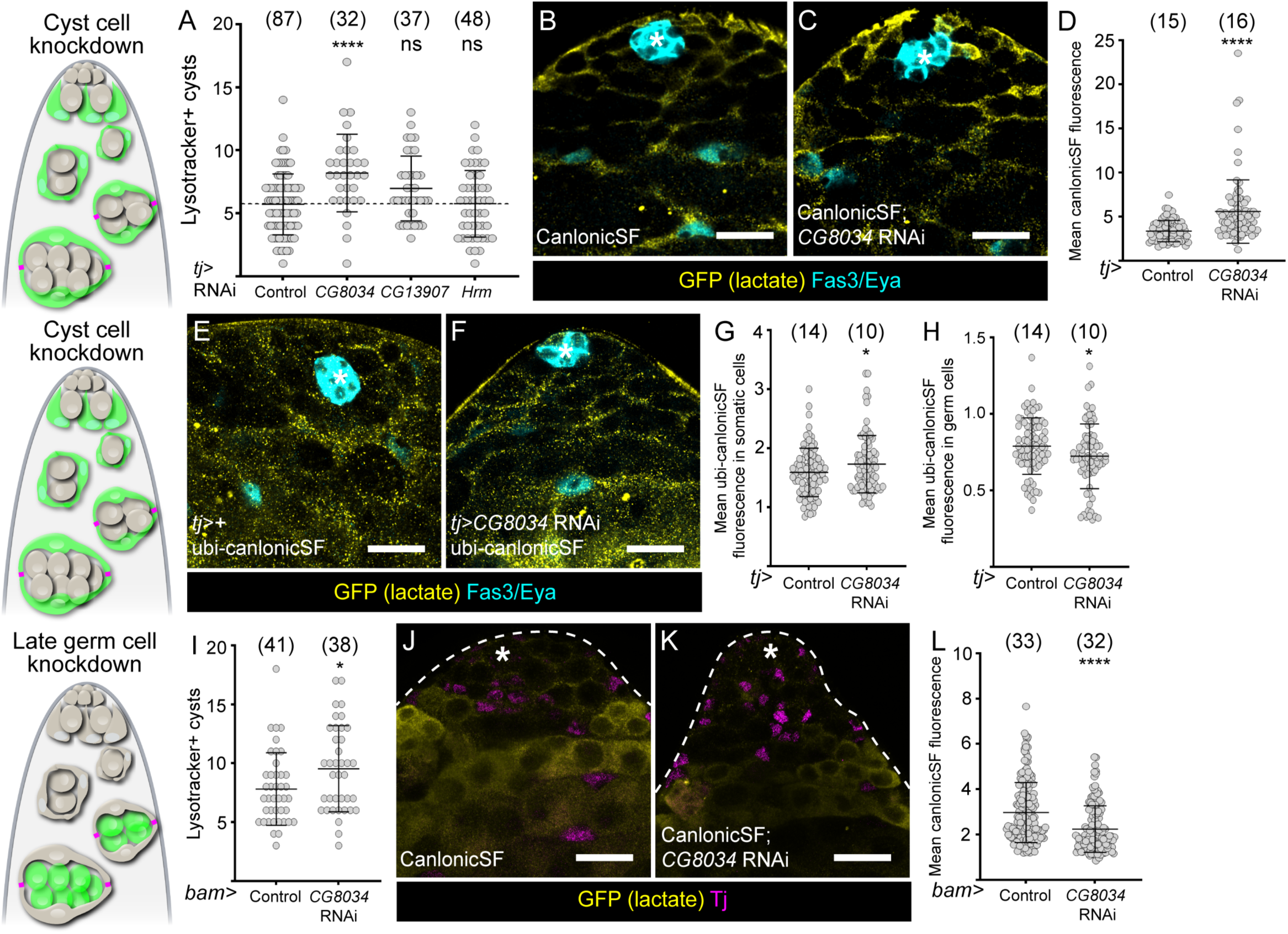
*CG8034/milkman* is required for lactate shuttling between the soma and the germline. (A) Graph showing the number of Lysotracker-positive germ cell cysts in control testes and testes in which *CG8034*, *CG13907* or *hrm,* encoding monocarboxylate transporters, were knocked down in cyst cells. **** denotes P < 0.0001, Kruskal Wallis and Dunn’s multiple comparisons tests. (B,C) Confocal images of testes in which the lactate reporter CanlonicSF (yellow) was expressed in cyst cells with *tj-Gal4* in control testes (B) or together with *CG8034* knockdown (C). Fas3 and Eya (cyan) label the hub and cyst cells, respectively. (D) Graph showing the mean intensity of CanlonicSF fluorescence in cyst cells in control testes and testes in which *CG8034* was knocked down in cyst cells. **** denotes P < 0.0001, student’s t test. (E,F) Confocal images of testes in which the lactate reporter CanlonicSF (yellow) was expressed ubiquitously in control testes (E) or in testes in which *CG8034* was knocked down in cyst cells with *tj-Gal4* (F). Fas3 and Eya (cyan) label the hub and cyst cells, respectively. (G) Graph showing the mean intensity of CanlonicSF fluorescence in cyst cells in control testes and testes in which *CG8034* was knocked down in cyst cells. * denotes P < 0.05, student’s t test. (H) Graph showing the mean intensity of CanlonicSF fluorescence in germ cells in control testes and testes in which *CG8034* was knocked down in cyst cells. * denotes P < 0.05, student’s t test (I) Graph showing the number of Lysotracker-positive cysts in control testes and testes in which *CG8034* was knocked down in differentiating germ cells with *bam-Gal4*. (J,K) Confocal images of testes in which the lactate reporter CanlonicSF (yellow) was expressed in germ cells with *bam-Gal4* in control (J) or together with *CG8034* knockdown (K). Tj (magenta) labels CySCs and early CCs. (K) Graph showing the mean intensity of CanlonicSF fluorescence in germ cells in control and *CG8034* knockdowns. *** denotes P < 0.001, student’s t test. All graphs: N values are shown in brackets and refer to the number of testes analysed. Scale bars: 20 μm. Diagrams to the left represent testis apical tips and highlight the lineage where gene expression was manipulated in green.

Similarly, to determine which MCTs were mediating lactate import into germ cells, we examined SLC16 family member expression in germ cell transcriptional clusters corresponding to spermatogonia and early spermatocytes (Fig. S5B). *hermes (hrm)* was the most highly expressed MCT, but was previously shown to transport pyruvate and not lactate (Velentzas et al., 2018). Instead, we noted that *mlm* was also expressed in germ cells, so we tested whether knocking down *mlm* in germ cells affected their survival. Indeed, expressing *mlm* RNAi with *bam-Gal4* resulted in a significant increase in lysotracker-positive germ cell cysts (Fig. 4I). Consistently, *mlm* knockdown in germ cells led to an autonomous decrease in CanlonicSF fluorescence (Fig. 4J-L), suggesting that Mlm mediates lactate import into germ cells.

### Somatic cells compartmentalise their energetic metabolism to ensure lactate production

Since glycolysis in cyst cells is critical for germline survival, we asked how cyst cell metabolism was set up to satisfy both the needs of germ cells and of cyst cells themselves. In particular, we wondered whether the pyruvate produced by glycolysis was a limiting resource or whether it was used both for lactate production for germ cells and for autonomous energy production in cyst cell mitochondria (Fig. 3A). We reasoned that if glycolysis fuelled cyst cell mitochondrial metabolism, knockdown of glycolytic enzymes should result in reduced mitochondrial activity. We used the mitochondrial membrane potential-sensitive dye tetramethyl rhodamine, methyl ester (TMRM) as a readout of mitochondrial activity. Knockdown of either *Pfk* or *Pyk* had no effect on TMRM fluorescence intensity in cyst cells (Fig. S6A,B). This suggested that glycolysis-derived pyruvate does not measurably contribute to cyst cell mitochondrial metabolism. To test whether mitochondrial consumption of pyruvate in cyst cells would result in reduced lactate production and germ cell support, we increased mitochondrial import of pyruvate in cyst cells by over-expressing the Mitochondrial pyruvate carrier, Mpc1. This manipulation resulted in a significant increase in dying germ cell cysts (Fig. 5A), indicating that pyruvate levels in cyst cells are limiting and that autonomous mitochondrial usage comes at the expense of germ cell support.

**Figure 5.**
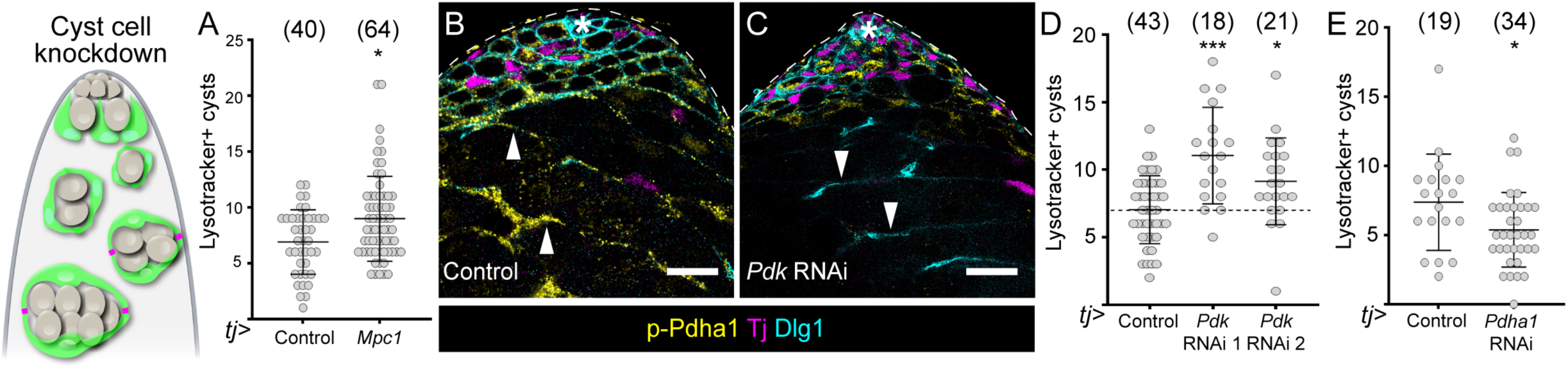
Compartmentalisation of cyst cell pyruvate metabolism ensures lactate production and germ cell support. (A) Graph showing the number of Lysotracker-positive germ cell cysts in control testes and testes in which *Mpc1* was over-expressed in cyst cells. * denotes P < 0.05, Mann-Whitney test. (B,C) Confocal images of a control testis (B) and a testis in which *Pdk* was knocked down in cyst cells with *tj-Gal4* (C), labelled with antibodies against phospho-Pdha1 (p-Pdha1, yellow), Tj to label CySCs and early cyst cells, and Dlg, to label cell outlines. Arrowheads highlight differentiated cyst cells where p-Pdha1 is visible in control but not upon *Pdk* knock down. The hub is indicated by an asterisk. (D) Graph showing the number of Lysotracker-positive cysts in control testes and testes in which *Pdk* was knocked down in cyst cells with *tj-Gal4*. *** denotes P < 0.001, * denotes P < 0.05, Kruskal Wallis and Dunn’s multiple comparisons tests. (E) Graph showing the number of Lysotracker-positive cysts in control testes and testes in which *Pdha1* was knocked down in cyst cells with *tj-Gal4*. * denotes P < 0.05, Mann-Whitney test. All graphs: N values are shown in brackets and refer to the number of testes analysed. Scale bars: 20 μm. The diagrams on the left represent a testis apical tip and highlights the lineage where gene expression was manipulated in green.

We then asked what normally ensures that pyruvate usage is steered towards lactate production. One possibility is that Mpc1 itself regulates flux; however, knockdown of *Mpc1* in cyst cells had no effect on germ cell death (Fig S6C). Therefore, we focused on regulation of the mitochondrial Pyruvate dehydrogenase complex (Pdh), a 3-enzyme (E1-3) complex which decarboxylates pyruvate to produce acetyl-coA. Pdh can be phosphorylated by Pyruvate dehydrogenase kinase (Pdk), which inhibits its activity (Fig. 3A). Using an antibody against phosphorylated Pdh, we observed staining in cyst cells in control testes (Fig. 5B), indicating that Pdk is active in cyst cells. Indeed, knockdown of *Pdk* using *tj-Gal4* resulted in loss of somatic phospho-Pdh staining (Fig. 5C), suggesting that Pdk normally acts to restrain Pdh activity, and therefore pyruvate consumption, in cyst cells. Consistently, knocking down *Pdk* resulted in significantly increased numbers of Lysotracker-positive dying germ cell cysts (Fig. 5D). Thus, similar to the over-expression of Mpc1, manipulations that increase autonomous pyruvate consumption in cyst cells result in decreased germ cell survival. Finally, we tested the effect of knocking down the gene encoding the α subunit of PdhE1, *Pdha1*, in cyst cells, hypothesizing that preventing mitochondrial consumption would increase the flux of pyruvate towards lactate production. Strikingly, *Pdha1* knockdown in cyst cells resulted in significantly fewer Lysotracker-positive germ cell cysts compared to control testes (Fig. 5E), indicating that preventing autonomous pyruvate usage in cyst cells is beneficial for germ cell survival. Together, these results show that cyst cell metabolism does not rely on pyruvate, and that phosphorylation of Pdh by Pdk is a key regulatory step that ensures that most pyruvate produced by glycolysis in cyst cells is steered towards the production of lactate to support the germline.

## Discussion

Male germ cells develop inside a specialised microenvironment as a result of a permeability barrier formed by gonadal somatic cells. How germ cell metabolism is sustained without access to circulating nutrients was unknown. Here we used metabolite sensors and targeted manipulations in germline and somatic cells to show that somatic gonadal cells promote germ cell survival by shuttling lactate.

We show that differentiating cyst cells upregulate glycolytic enzymes, to produce lactate which is transferred to germ cells. In germ cells, lactate consumption through conversion into pyruvate is essential for viability, demonstrating that one of the key functions of cyst cells is to support germline metabolism by providing trophic metabolites. Finally, our results show that manipulating cyst cell metabolism to increase or decrease autonomous pyruvate consumption in mitochondria results in converse changes in germ cell survival, emphasising the importance of ensuring proper resource allocation in cyst cells: glycolysis and its product lactate are dedicated to the support of germ cells. While direct *in vivo* evidence is lacking, several pieces of evidence suggest that lactate shuttling between somatic and germline cells is conserved. In mammals, Sertoli cells establish the blood-testis barrier, enclosing spermatocytes in the lumen of the testis tubules, away from the blood vessels (Dym and Fawcett, 1970; Stanton, 2016). Although the testis as a whole depends on glucose metabolism, experiments using cultured cells suggest that germ cells themselves do not metabolise glucose, while Sertoli cells do (Boussouar and Benahmed, 2004; Jutte et al., 1981; Jutte et al., 1982; Mita and Hall, 1982; Robinson and Fritz, 1981). Thus it seems likely that *in vivo*, Sertoli cells break down carbohydrates to produce lactate, which is secreted and used in germ cells for energy metabolism, similar to the compartmentalization of carbohydrate metabolism between neurons and glia (Brooks, 2018; Gladden, 2004).

### Metabolic compartmentalization in somatic cells of the testis

Our findings highlight that cyst cells compartmentalize their metabolism. Despite a strong upregulation of glycolytic enzymes, glycolysis is not autonomously required in cyst cells and does not appreciably contribute to cyst cell fate or to their mitochondrial activity. This compartmentalisation appears to be conserved with mammals, as *in vitro* work indicates that although cultured Sertoli cells take up and metabolise glucose, only a small fraction of this glucose is consumed through oxidation in mitochondria (Grootegoed et al., 1986; Robinson and Fritz, 1981). Pdk plays a key role in cyst cell compartmentalisation but in cultured Sertoli cells treatment with a Pdk inhibitor prevents increases in lactate production upon stimulation with Fibroblast Growth Factor (FGF), but does not reduce basal lactate production (Regueira et al., 2015). Thus, lactate secretion by Sertoli cells may be both intrinsically and extrinsically regulated in mammals. Indeed, in Sertoli cells, the pituitary-derived follicle stimulating hormone and FGF, secreted by the germline, regulate expression of glycolytic enzymes and glucose transporters, as well as expression and activity of Ldh, linking upregulation of lactate production to puberty (Galardo et al., 2008; Mita et al., 1982; Regueira et al., 2015; Riera et al., 2002). In Drosophila cyst cells, it remains unknown what regulates Pdk expression and/or activity.

Notably, in previous work, we showed that cyst cell differentiation involves changes in mitochondrial morphology and activity, suggestive of increased metabolic requirements with differentiation (Sainz de la Maza et al., 2022). Thus, multiple metabolic pathways change coordinately during cyst cell development, to allow for both autonomous needs in differentiation, and for non-autonomous support of the germline. Our results indicate clearly that glycolysis does not contribute to autonomous functions, and suggest that other, as yet unknown, metabolic pathways are upregulated to promote cyst cell differentiation autonomously. Thus, cyst cell metabolism is compartmentalised such that distinct metabolic pathways contribute to distinct functions in cyst cells, ensuring that sufficient resources are allocated both to autonomous needs and to non-autonomous support of the germline.

### Lactate in the germline

Our data indicate that in the Drosophila testis, spermatogonia become dependent on lactate for survival. Indeed, preventing conversion of lactate into pyruvate in germ cells results in increased germ cell death, implying that germ cells consume lactate. Strikingly, the stage-specific dependence of male germline cells on lactate appears conserved in mammals: spermatocytes and round spermatids specifically depend on exogenous lactate when in culture and degenerate in its absence (Jutte et al., 1981; Mita and Hall, 1982). While best studied in rats, evidence also indicates that lactate suppresses apoptosis in cultured human testis sections (Erkkila et al., 2002). The reasons for this lactate dependence in developing germ cells are still not fully understood, but in its absence, RNA and protein synthesis decreased, as did ATP levels, implicating lactate in energy production via mitochondrial oxidation (Grootegoed et al., 1984; Jutte et al., 1981; Mita and Hall, 1982). One possibility is that the activity of Ldh tends to favour lactate production, resulting in the depletion of pyruvate in germ cells, such that high levels of lactate are required to allow germ cells to use pyruvate autonomously (Grootegoed et al., 1984). Intriguingly, in both flies and mammals, the lactate dependence of germ cells coincides with their isolation behind the barrier formed by somatic gonadal cells (Fairchild et al., 2015; Stanton, 2016). Thus, it is possible that one reason for isolating germ cells is to create a tightly controlled environment in which lactate levels can be elevated locally, enabling pyruvate consumption in the germline.

### Lactate as a key currency in intercellular metabolic exchange

Nonetheless, it is notable that cyst cells provide lactate and not another metabolite such as glucose or pyruvate to the germline. In addition to testes, metabolic exchange between cells in the same tissue has been described in several contexts. In particular, astrocytes provide lactate to neurons in flies and mammals, as do Paneth cells to intestinal stem cells in the mammalian gut (Brooks, 2018; Rodriguez-Colman et al., 2017; Volkenhoff et al., 2015). Thus, support cells in many different tissues appear to use lactate as a metabolic currency, suggesting that lactate, and therefore mitochondrial oxidation, is a preferred energy source in cells that depend on others for support. This dependence on lactate as a nutrient is especially striking in the case of both neurons and germ cells, two cell types that are long-lived and which need to protect their DNA from reactive oxygen species. Current models often suggest that proliferating cells depend on glycolysis to provide precursors for nucleotide synthesis while shunning oxidative phosphorylation to prevent DNA damage (Vander Heiden et al., 2009). However, in the Drosophila testis, spermatogonia are still proliferating when the surrounding cyst cells form tight junctions, such that the dependence on cyst cell-derived lactate does not correspond to the end of mitotic stages. Instead, one possibility is that since lactate conversion into pyruvate generates NADH, the key reason for reliance on lactate may be to regulate cellular redox, which would in turn have critical effects on energy generation, as well as macromolecule synthesis (Hosios and Vander Heiden, 2018).

Overall, our work sheds light on a key function of somatic support cells and raises the possibility that somatic metabolism may contribute to male infertility in humans. Our finding that lactate sustains germ cell metabolism will have important implications for the future development of *in vitro* culture protocols. Understanding not only germ cell metabolism, but the nature of the support provided by somatic gonadal cells will enable improved germ cell culture protocols which currently involve co-culture with Sertoli cells (Ishikura et al., 2021). By understanding how somatic cells support the germline, it may be possible to replace somatic cells in co-culture with liquid media, a crucial step towards clinical applications of *in vitro* gametogenesis for the treatment of infertility.

## Materials and methods

### Fly stocks and husbandry

Crosses were raised at room temperature. Males were collected 0-3 days after eclosion and shifted to 29°C for 10 days. For clone generation, crosses were raised at 25°C, adult males were collected 0-3 days after eclosion and heat shocked at 37°C in a water bath for 1 hour.

We used Flybase (release FB2025_02) to find information on stocks, gene groups and pathways (Ozturk-Colak et al., 2024). Stocks obtained from the Bloomington Drosophila Stock Center (BDSC, NIH P40OD018537), the Kyoto Drosophila Stock Center at Kyoto Institute of Technology (DGRC) and the Vienna Drosophila Resource Center (VDRC, www.vdrc.at) were used in this study. The following stocks were used:

*Tret1-GFPnls* (BDSC #94540), *Aldo-GFP* (DGRC #115279), *PyK-HA* (gift from S. Schirmeier), *Tj-Gal4*, *Hh-Gal80* (Herrera et al., 2021), *nos-Gal4::VP16* (Amoyel Lab), *bam-Gal4::VP16*, *Tret1 RNAi* (VDRC #8126 and BDSC #42880), *Treh RNAi* (VDRC #30730), *Pgi RNAi* (VDRC #24257), *Pfk RNAi* (VDRC #3016), *Gapdh1 RNAi* (VDRC #100596), *Pgk RNAi* (VDRC #33797), *Pgam1 RNAi* (VDRC #52336), *Eno RNAi* (VDRC #110090 and VDRC #330201), *PyK RNAi* (VDRC #49533), *UAS-mCherry-nls* (BDSC #38424), *UAS-RFP* (BDSC #30556), *UAS-CanlonicSF* (BDSC #94537), *Ldh RNAi* (BDSC #33640), *UAS-PyronicSF* (BDSC #94533 and BDSC #94535), *UAS-Diap1* (Gift from N. Tapon), *CG8034 RNAi* (BSDC #32340), *CG13907 RNAi* (VDRC #107339), *Hrm RNAi* (VDRC #7314), *UAS-MPC1* (BDSC #83687), *Pdk RNAi* (BDSC #28635 and BDSC #35142), *Pdha1 RNAi* (VDRC #40410), *MPC1 RNAi* (VDRC #15858 and VDRC #103829), *y,w,hsflp^122^,Tub>Gal4,UAS-nlsGFP;; FRT^82B^,Tub>Gal80*, *FRT^82B^,ry^506^*, *FRT^82B^,ald1^EY13155^* (gift from M. Simonelig).

*Ubi-CanlonicSF* was generated by DNA synthesis (ThermoFisher Scientific) of the sequence for CanlonicSF (Aburto et al., 2022) and subcloning into pUbi-AttB (gift from N. Tapon). The resulting plasmid was injected by Bestgene Inc into embryos carrying attP sites VK00005 and VK00037 (BDSC #9725 and 9752) and adult flies were screened for integration using eye colour to establish stable lines.

### Single-nucleus RNA-seq data analysis

We utilized the publicly available, single-nucleus transcriptome of Drosophila testes from Li et al, 2022, using the clusters and annotations generated by Raz *et al*, 2023. Plots were generated either using the online platform ASAP (Gardeux et al, 2017) or the Seurat package (Hao *et al*, 2023) in R Studio (Posit Team, 2024).

### Immunohistochemistry and cellular dyes

Dissected abdomens were fixed in 4% paraformaldehyde in PBS for 15 minutes. Samples were washed twice in PBS, 0.5% Triton X-100 for 30 minutes then blocked in PBS, 1% BSA, 0.2% Triton X-100 (PBTB) for one hour, before overnight incubation in primary antibodies diluted in PBTB. Samples were then washed twice in PBTB for 30 minutes, and incubated in secondary antibodies diluted in PBTB for 2 hours at room temperature, then washed in PBS, 0.2% Triton X-100, and mounted on slides with Vectashield mounting medium for imaging. For Lysotracker staining, abdomens were incubated in Schneider’s insect medium (Sigma, S0146) supplemented with Lysotracker Red DND-99 (1:1000, Thermo Fisher, L7528) for 1 hour prior to fixation and staining as above. For p-Pdha staining, flies were dissected in 10mM TrisHCl pH6.8, 0.18M KCl, with phosphatase inhibitors (50mM NaF, 10mM NaVO4, 10mM beta-glycerophosphate), followed by standard fixation and staining. For EdU staining, abdomens were incubated in Schneider’s medium with 10μM EdU (Abcam, ab146186) for 30 minutes prior to fixation. Following secondary antibody incubation, fluorescent azides were conjugated to EdU by click reaction for 30 minutes in PBS with 2.5 μM Alexa picolyl azide (Click Chemistry Tools), 0.1□mM THPTA, 2□mM sodium ascorbate and 1□mM CuSO4. For measurement of mitochondrial potential, dissected abdomens from *tj-Gal4, UAS-CD8GFP* flies were incubated in Schneider’s insect medium with 25nM TMRM (ThermoFisher Scientific #T668) for 30 minutes before being mounted and imaged live.

The following primary antibodies were used: chicken anti-GFP (1:500, Aves Labs, GFP-1010), rabbit anti-GFP (1:500, Thermo Fisher, A6455),), rabbit anti-Zfh1 (1:5000, this study), rabbit anti-HA (1:200, Cell Signaling, C29F4), guinea pig anti-Tj (1:5000, gift from D. Godt), guinea-pig anti-Zfh1 (1:3000 (Wang et al., 2025)), rabbit anti-Dcp1 (1:100, Cell Signaling, 9578), rabbit anti-phospho Pdha (1:100, Abcam, ab92696). Mouse anti-Eya (eya10H6, 1:20, deposited by S. Benzer/N.M. Bonini), mouse anti-Fas3 (7G10, 1:20, deposited by C. Goodman), rat anti-CadN (1:20), rat anti-De-cad (1:20), rat anti-Vasa (1:20, deposited by A.C. Spradling/D. Williams) and mouse anti-Dlg (4F3, 1:20, deposited by C. Goodman) were obtained from the Developmental Studies Hybridoma Bank created by the NICHD of the NIH and maintained at The University of Iowa.

The rabbit anti-Zfh1 antibody was generated by GenScript using the same recombinant antigen (amino acids 648–775 of Zfh1 isoform PB) previously used for generating the guinea pig antibody (Wang et al., 2025). This antigen was injected into two rabbits. The resulting serum was purified by antigen affinity column to obtain concentrated antiserum.

### Mass spectrometry for metabolite detection

Testes from *tj-Gal4; UAS-CD8-GFP* flies were dissociated as described in Sainz de la Maza *et al*, 2022 to generate a cell suspension. GFP^+^ somatic cells and GFP^-^germ cells were sorted using a BD FACSAria Fusion Cell Sorter. Cells were spun down at 1550 RPM for 10 minutes at 4°C, resuspended in 1:3:1 chloroform/methanol/water for lysis and incubated for 1h at 4°C with rocking. Lysates were spun down at 13,000 RPM for 3 minutes at 4°C and supernatants were extracted and stored at −80°C.

Mass spectrometry analysis of the samples was carried out at the Polyomics Facility at the University of Glasgow. Hydrophilic interaction liquid chromatography (HILIC) was carried out on a Dionex UltiMate 3000 RSLC system (Thermo Fisher Scientific) using a ZIC-pHILIC column (150 mm × 4.6 mm, 5 μm column, Merck Sequant). Samples were eluted with a linear gradient of 20 mM ammonium carbonate in water and acetonitrile. For the MS analysis, a Thermo Orbitrap QExactive (Thermo Fisher Scientific) in polarity switching mode was used. To confirm the identity of metabolites, standards were run alongside the samples to match signals based on accurate mass and retention time. Signals that did not match authentic standards were annotated based on accurate mass. Identification of peaks and quantification of metabolite abundance were obtained using the web-based tool PiMP (Gloaguen et al., 2017). For PCA analysis and plotting of the annotated peaks, the data produced by PiMP was uploaded to Metaboanalyst 6.0 (Pang et al., 2024).

### Image acquisition and analysis

Images were acquired using Zeiss LSM800 and LSM880 confocal microscopes and analysis and quantifications were performed with Fiji (Schindelin et al., 2012). For TMRM experiments, images were acquired on a Zeiss LSM980 using Airyscan detectors. Imaris 9.9 (Oxford Instruments) software was used to reconstruct the three-dimensional structure of the entire somatic network and measure the mean fluorescence of TMRM within the soma.

Clonal CySCs were identified as Zfh1-positive cells adjacent to the hub that were also positive for GFP, while clonal cyst cells were identified as GFP-positive, Eya-positive cells. To quantify clone-associated cell death, all GFP-positive, Eya-positive cyst cells were assessed for the presence of a Lysotracker-positive germ cell cyst immediately adjacent; the fraction of cyst cells with a dying cyst was plotted.

For CySC counts, CySCs were identified as Zfh1-positive and Eya-negative cells.

For quantifications of metabolite reporters, control and experimental flies were dissected on the same day and processed simultaneously. To account for variability between samples, mean fluorescence intensity was normalised. When reporters were expressed in somatic cells, intensity values were normalised to a germ cyst, where the reporter was not expressed. When the reporter was expressed in differentiating germ cells with *bam-Gal4*, values were normalised to the stem cell area. Finally, when the reporter was expressed ubiquituously, values were normalised to the hub.

Statistical tests were carried out using GraphPad Prism. Non-parametric ANOVA was used to compare numbers of lysotracker-positive germ cysts between experimental conditions; to compare fluorescence intensities we used ANOVA when more than two experimental groups were present and t-test when only two groups were tested. Clone recovery and clone-associated cell death rates were compared using Fisher’s exact test. Bars and whiskers on all graphs show mean ± SD.

## Acknowledgements

We are especially grateful to Gavin Blackburn from the University of Glasgow for invaluable help analysing the mass spectrometry data. We thank colleagues in the Drosophila community and the Bloomington, Vienna and Kyoto stock centres for fly stocks and reagents. We are grateful to many colleagues including Vilaiwan Fernandes, Alex Gould, Nazif Alic, Joaquina Delas, Roberto Mayor and members of the Amoyel and Fernandes labs for critical discussions and comments on the manuscript.

## Author contributions

Conceptualisation: MA, DSdlM; Investigation: DSdlM, HJ, CIB, SP; Supervision: MA; Funding acquisition: MA; Writing – original draft: MA, DSdlM; Writing – review and editing: MA, DSdlM.

## Data and materials availability

All data are available in the manuscript or the supplementary materials. All Drosophila stocks generated in the course of this study are available from the authors and have been deposited at the Bloomington Drosophila Stock Center where appropriate.

---

**Figure S1. Somatic and germ cells have different metabolic profiles**

(A) UMAP representation showing the clustering and annotation generated by Raz et al of the Fly Cell Atlas single-nucleus RNA sequencing of the testis. (B) Gene expression plot showing *tj* expression on the testis UMAP identifying clusters 62, 36 and 58 as corresponding to CySCs and early cyst cells. (C) Gene expression plot showing *vasa (vas)* expression on the testis UMAP representation, identifying clusters 25, 22 and 5 as corresponding to early stages of germ cell development. (D) Table listing the two most significantly enriched GO terms in genes enriched in early cyst cells (clusters 62, 36 and 58) compared to early germline clusters (25 and 22). The genes contained within the GO categories are shown, and those encoding glycolytic enzymes are highlighted in yellow. (E) Principal component analysis score plot from the mass spectrometry data from the independent replicates of sorted cyst (blue dots) and germ cells (grey dots). (F) Volcano plot showing the relative enrichment of all annotated mass spectrometry peaks in cyst cells compared to germ cells.

**Figure S2. Knockdown of genes encoding glycolytic enzymes does not affect CySC numbers or cyst cell survival.**

(A) Confocal image of a testis expressing GFP (yellow) in somatic cells driven by *tj-Gal4*. Vasa (white) labels germ cells, DE-Cad (cyan) labels cell outlines. Scale bar: 20 μm. The diagram on the left represents a testis apical tip and highlights *tj-Gal4* expression in green. (B) Graph showing the number of Zfh1-positive, Eya-negative cells in control testes or testes in which the indicated genes encoding glycolytic enzymes were knocked down in cyst cells. Significance was assessed using Kruskal Wallis and Dunn’s multiple comparisons tests. (C) Graph showing the number of Dcp-1-positive cells in control testes or testes in which the indicated genes encoding glycolytic enzymes were knocked down in cyst cells. Significance was assessed using Kruskal Wallis and Dunn’s multiple comparisons tests. (D) Graph showing the recovery rate of control and A*ldo^EY13155^* somatic clones. No significant difference was detected using Fisher’s exact test. All graphs: N values are shown in brackets and refer to the number of testes analysed.

**Figure S3. Driver expression and RNAi knockdown in the germline.**

(A,B) Confocal images of testes expressing mCherry (yellow) in early germ cells cells driven by *nos-Gal4* (A) or RFP (yellow) in differentiating germ cells driven by *bam-Gal4* (B). Vasa (white) labels germ cells, Zfh1 (magenta) labels CySCs, Fas3 and Eya (cyan) label the hub and cyst cells, respectively. Scale bar: 20 μm. The diagrams on the left represent a testis apical tip and highlight the domains of *nos-Gal4* and *bam-Gal4* expression in green. (C,D) Confocal images of testes from *Pyk-HA* control flies (C) or flies in which *Pyk* was knocked down in early germ cells with *nos-Gal4*, labelled with antibodies against HA (yellow) and Fas3 and Eya (cyan) to label the hub and cyst cells respectively. Scale bars: 20 μm.

**Figure S4. *Ldh* is not required autonomously in CySCs or cyst cells.**

(A) Graph showing the number of Zfh1-positive, Eya-negative cells in control testes or testes in which *Ldh* was knocked down in cyst cells. Significance was assessed using a Mann-Whitney test. (B) S-phase index of Zfh1-positive CySCs in control testes and testes in which *Ldh* was knocked down in cyst cells. Significance was assessed using a student’s t test. (C) Graph showing the number of Dcp-1-positive cells in control testes and testes in which *Ldh* was knocked down in cyst cells. Significance was assessed using a Mann-Whitney test. (D) Graph showing the number of Lysotracker-positive germ cell cysts in control testes and testes in which *Ldh* was knocked down in cyst cells, with or without overexpression of the apoptosis inhibitor Diap. *** denotes P < 0.001, **** denotes P < 0.0001, determined by Kruskal Wallis and Dunn’s multiple comparisons tests. All graphs: N values are shown in brackets and refer to the number of testes analysed.

**Figure S5. Expression of monocarboxylate transporters in somatic and germ lineages.**

(A) Dot plot showing average expression from the Fly Cell Atlas dataset of the indicated genes encoding MCTs in the cell clusters corresponding to CySCs (62) and early cyst cells (36 and 58). (B) Dot plot showing average expression from the Fly Cell Atlas dataset of the indicated genes encoding MCTs in the cell clusters corresponding to early (25), and late spermatogonia (22) and cells at the spermatogonia-spermatocyte transition (5). The size of the dots indicates the percent of expressing cells in each cluster, and the colour represents the expression level. Note that *chaski* was not identified in this dataset.

**Figure S6. Mitochondrial activity in cyst cells does not depend on autonomous pyruvate consumption.**

(A) Airyscan image of a testis in which GFP (cyan) was expressed with *tj-*Gal4 to identify cyst cells and labelled with the mitochondrial membrane potential-sensitive dye TMRM (yellow). Scale bar: 20 μm. (B) Graph showing TMRM mean intensity in cyst cells in control testes and testes in which *Pfk* or *Pyk* were knocked down. Significance was determined by Kruskal Wallis and Dunn’s multiple comparisons tests. (C) Graph showing the number of Lysotracker-positive cysts in control testes and testes in which *Mpc1* was knocked down in cyst cells. Significance was determined by Kruskal Wallis and Dunn’s multiple comparisons tests. All graphs: N values are shown in brackets and refer to the number of testes analysed.

