## Supplementary material for "Somatic cells compartmentalise their metabolism to sustain germ cell survival": Table S1

**Table S1: RNAi lines used and viability when crossed to *tubulin-Gal4***

Table showing the genes encoding glycolytic enzymes targeted, stock reference numbers and observed viability (relative to expected) when crossed to *tubulin-Gal4/TM6B.*

| **Gene** | **RNAi stock** | **% viability**  **(N)** | **Used in experiments** |
| --- | --- | --- | --- |
| Control | - | 104.2  (119) | N/A |
| *treh* | VDRC #30730 | 0  (205) | Y |
| *hex-A* | VDRC #21054 | 102  (505) | N |
| *hex-C* | VDRC#35337 | 112  (288) | N |
| *pgi* | VDRC #24257 | 0  (117) | Y |
| *pfk* | VDRC #3016 | 15  (196) | Y |
| *aldo* | VDRC #27541 | 81  (42) | N |
| *tpi* | VDRC #25643 | 143  (245) | N |
| *gapdh1* | VDRC #100596 | 3  (142) | Y |
| *gapdh1* | VDRC #31632 | 73  (379) | N |
| *gapdh2* | VDRC #50351 | 89  (192) | N |
| *pgk* | VDRC #33797 | 40*  (269) | Y |
| *pgam1* | VDRC #52336 | 0  (137) | Y |
| *eno* | VDRC #330201 | 0  (201) | Y |
| *eno* | VDRC #110090 | 0  (265) | Y |
| *pyk* | VDRC #49533 | 0  (170) | Y |
| *pyk* | VDRC #35165 | 107  (129) | N |

* For *pgk* RNAi, females were recovered at 69% of expected rates (N = 157), but no males were recovered (0% viability, N = 112).
