## Supplementary material for "Somatic cells compartmentalise their metabolism to sustain germ cell survival": Fig. S1

A

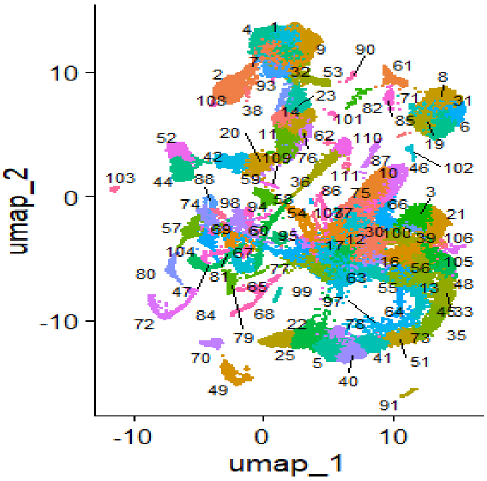


B

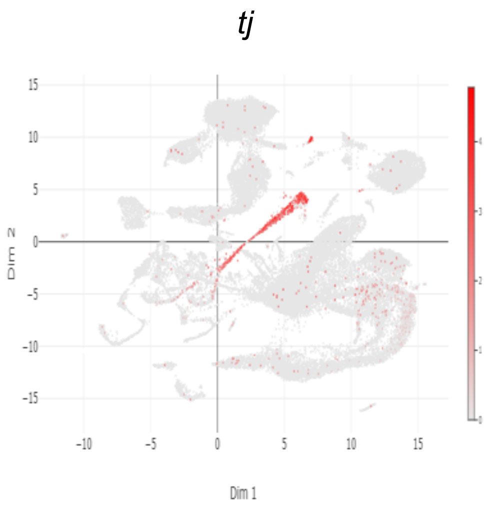


C

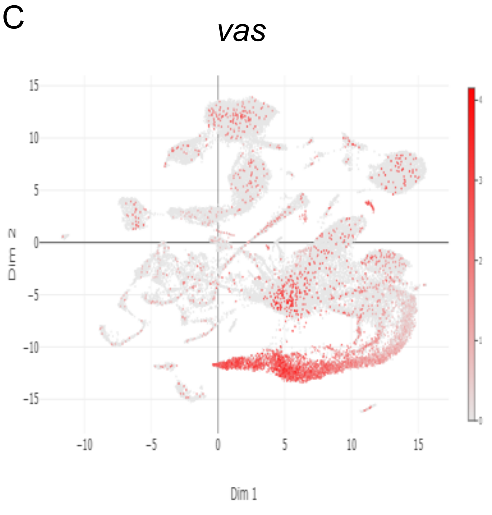


D

| Comparison of somatic clusters 62, 36, 58 vs germ clusters 25 and 22 | | | | | | | | | | |
|---|---|---|---|---|---|---|---|---|---|---|
| GO term (FDR value) | Genes | | | | | | | | | |
| Biosynthesis of amino acids (FDR=0,003) | *Eno* | *Gapdh2* | *Pfk* | *Pgam1* | *Tpi* | *aay* | *Alat* | *Arg* | *Bcat* | *CG3483* |
| | *CG32026* | *Glts* | *Gs1* | *Gs2* | *Idh* | *Idh3g* | *Irp-1B* | *P5CS* | *Pcb* | *Phgdh* |
| | *Psat* | *Pykl2* | *Pykl5* | *Sams* | *Taldo* | *Tkt* | *Tktl* | | | |
| Carbon metabolism (FDR=0,003) | *Eno* | *Gapdh2* | *Hex-A* | *Pfk* | *Pgam1* | *Tpi* | *aay* | *Acat1* | *Acat2* | *AcCoAS* |
| | *Alat* | *CG3483* | *CG5577* | *CG17544* | *CG17896* | *CG32026* | *CG32487* | *CG32488* | *Fbp* | *G6pd* |
| | *G6pdl* | *Gdh* | *Hibadh* | *Idh* | *Idh3g* | *Irp-1B* | *Mdh1* | *Mdh2* | *Ogdh1* | *Pcb* |
| | *Pgd* | *Phgdh* | *Psat* | *Pykl2* | *Pykl5* | *SdhA* | *SdhAL* | *Taldo* | *Tkt* | *Tktl* |

E

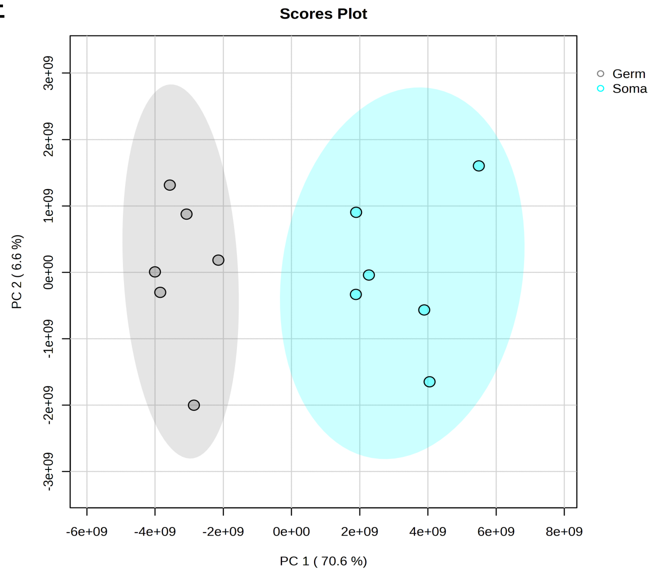


F

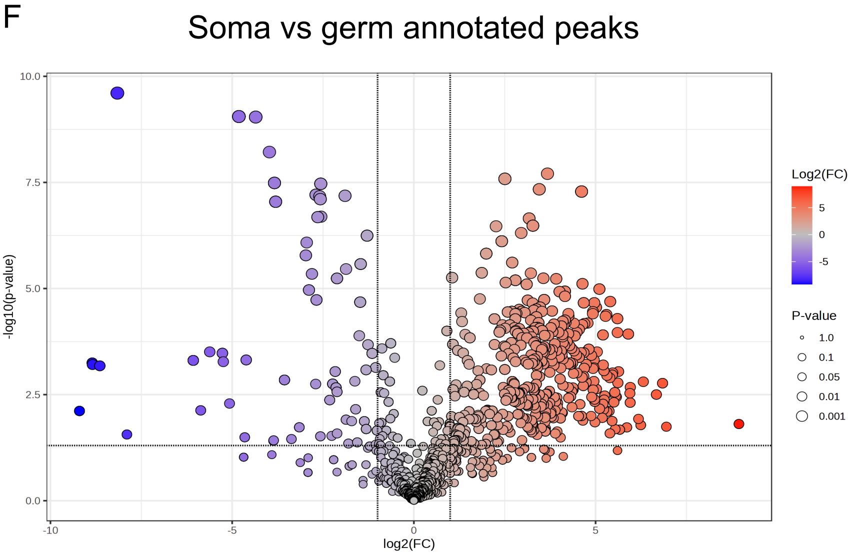
