## Supplementary figures and images for "Somatic cells compartmentalise their metabolism to sustain germ cell survival"

### Fig. S2

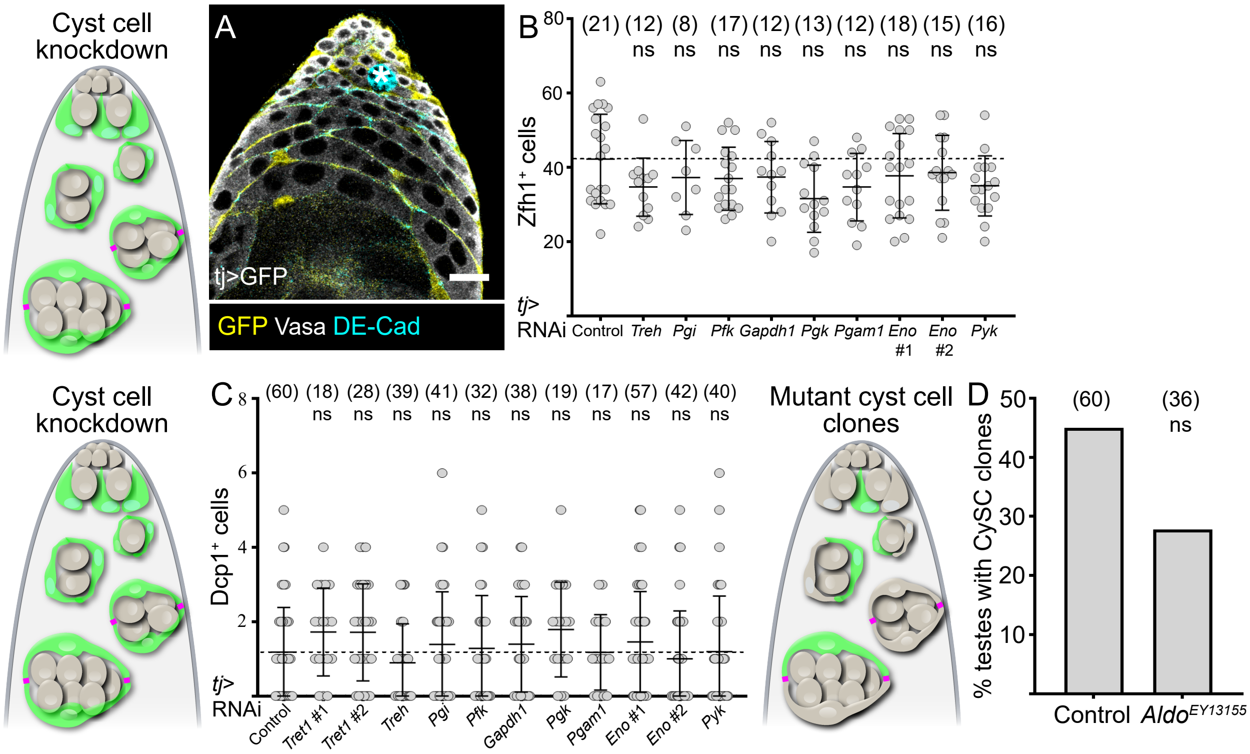

### Fig. S3

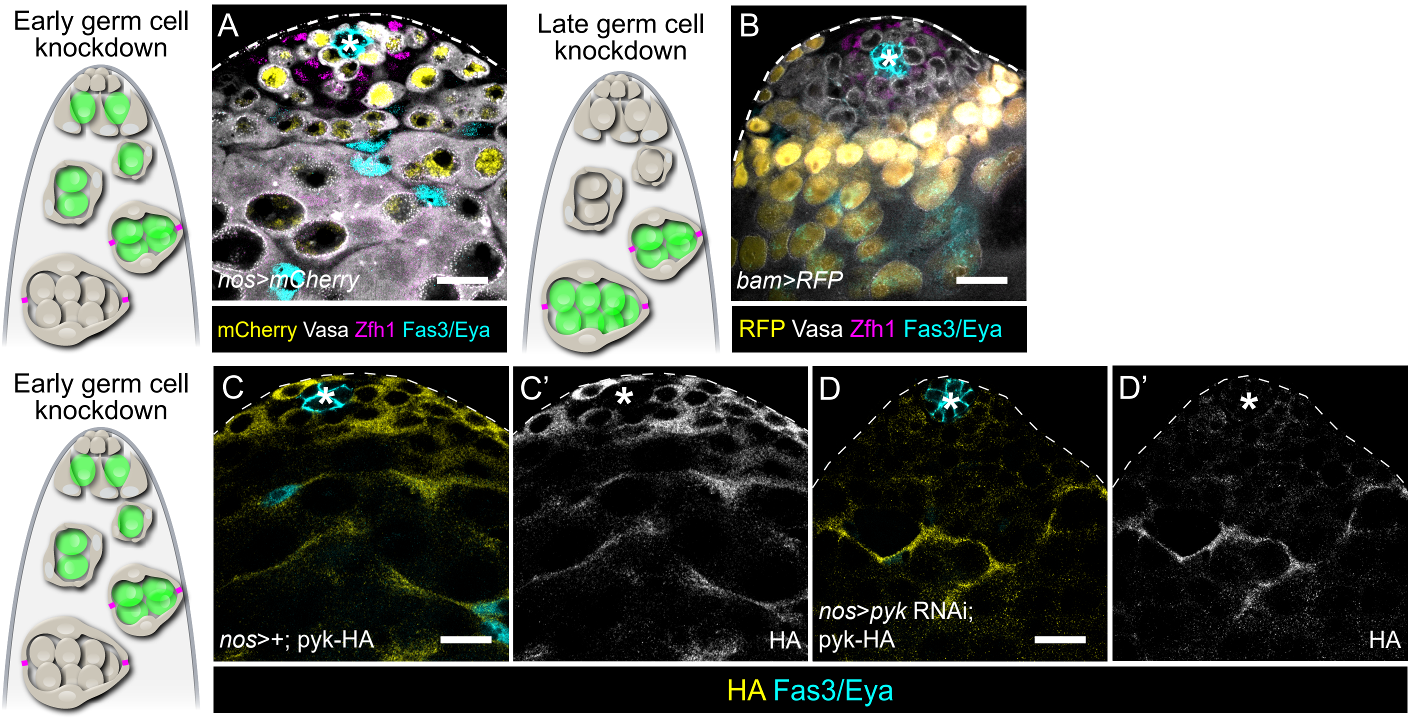

### Fig. S4

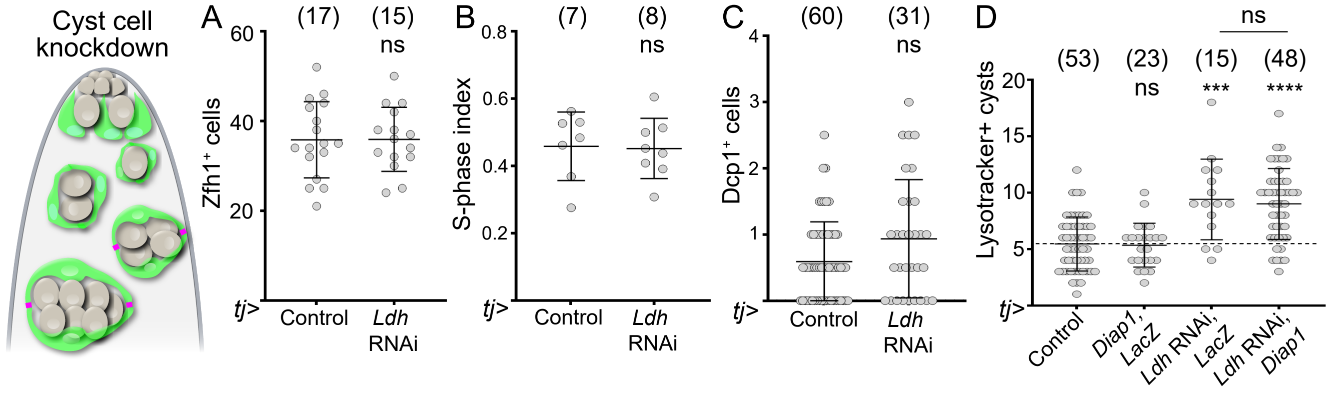

### Fig. S5

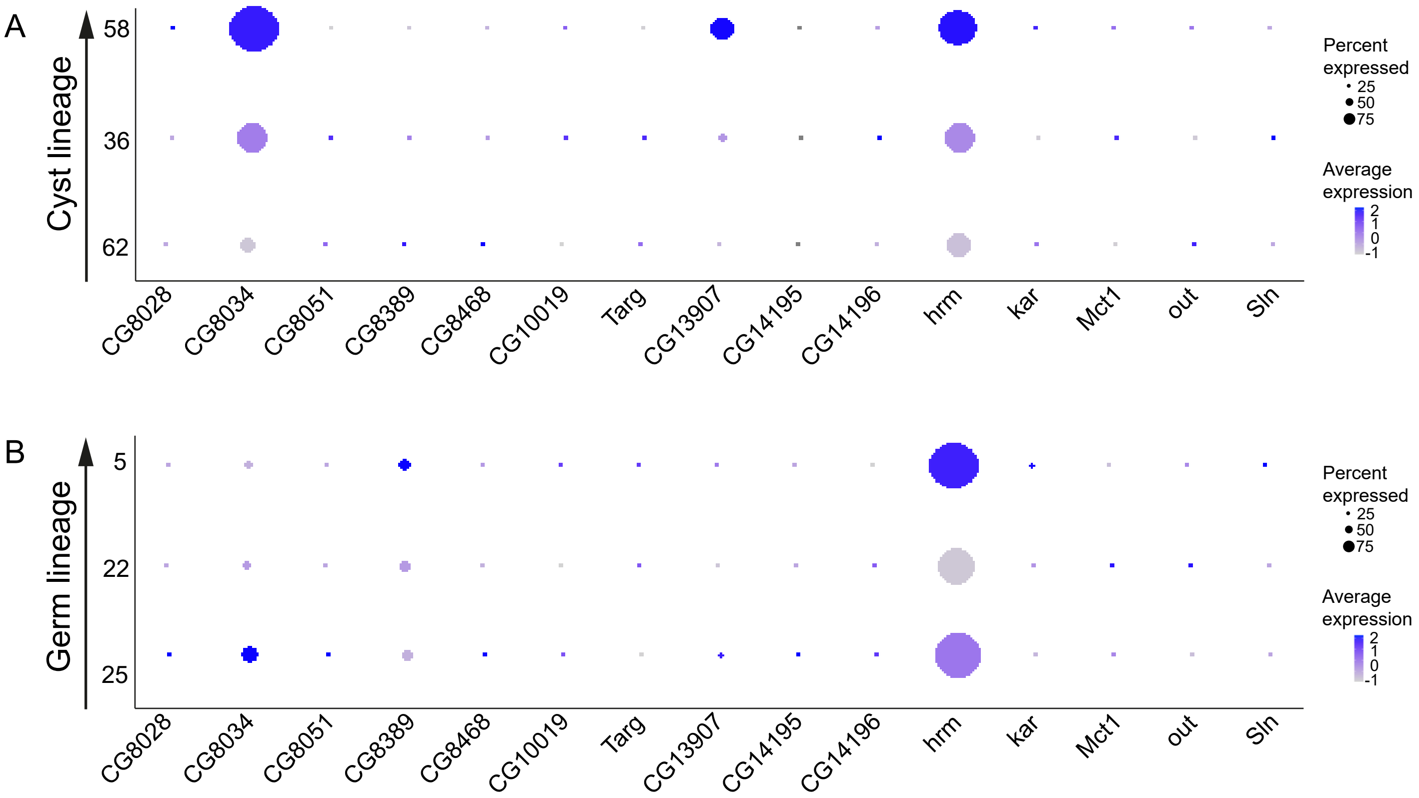

### Fig. S6

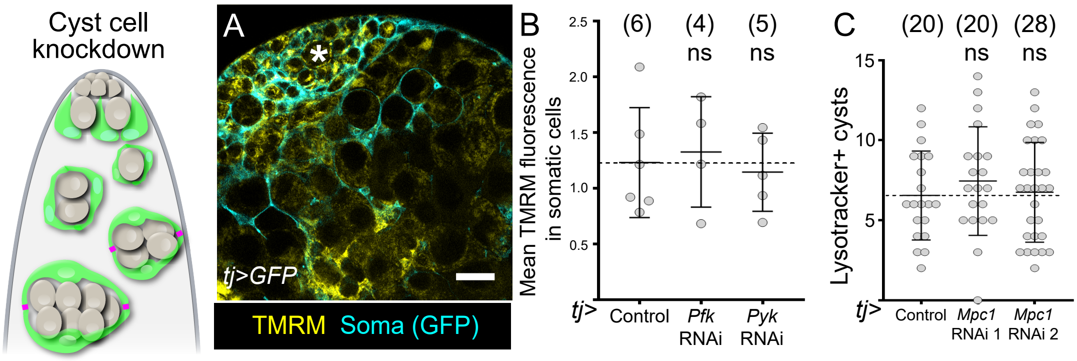
